# Partial Mimicry of the Microtubule Binding of Tau by Its Membrane Binding

**DOI:** 10.1101/2022.11.06.515359

**Authors:** Matthew MacAinsh, Huan-Xiang Zhou

## Abstract

Tau, as typical of intrinsically disordered proteins (IDPs), binds to multiple targets including microtubules and acidic membranes. The latter two surfaces are both highly negatively charged, raising the prospect of mimicry in their binding by tau. The tau-microtubule complex was recently determined by cryo-EM. Here we used molecular dynamics simulations to characterize the dynamic binding of tau K19 to an acidic membrane. This IDP can be divided into three repeats, each containing an amphipathic helix. The three amphipathic helices, along with flanking residues, tether the protein to the membrane interface. The separation between and membrane positioning of the amphipathic helices in the simulations are validated by published EPR data. The membrane contact probabilities of individual residues in tau show both similarities to and distinctions from native contacts with microtubules. In particular, a Lys that is conserved among the repeats forms similar interactions with membranes and with microtubules, as does a conserved Val. This partial mimicry facilitates both the membrane anchoring of microtubules by tau and the transfer of tau from membranes to microtubules.

---

The microtubule associated protein tau exists in one of six isoforms, differing in the presence or absence of two adjacent N-terminal inserts and of the second of four partially repeated microtubule binding regions (MTBRs).^1^ During neuronal development, tau is enriched in the axon and is responsible for microtubule assembly and stabilization.^2, 3^ In addition, it regulates axonal transport by acting as a gate for dynein and kinesin motor proteins as they move along microtubules.^4^ Tau mediates these functions by binding directly to and diffusing along microtubules, through its four MTBRs (R1-R4). Microtubule-bound tau can form condensates known as “envelopes”.^5^ In solution, tau can also form condensates that facilitate further aggregation into neurofibrillary tangles.^6, 7^ Accumulation of tau tangles in the brain is a hallmark of multiple neurodegenerative diseases including Alzheimer’s.

The functions and disease linkage of tau are intimately related to its nature of being intrinsically disordered. Like many other intrinsically disordered proteins (IDPs), tau bind multiple targets. In addition to microtubules, the targets include annexins A2 and A6 (interacting with tau’s extreme N-terminus)^8^ and Src-family kinases such as Fyn (interacting with tau’s proline-rich region).^9^ Binding to such membrane-bound targets may contribute to the localization of tau at membrane surfaces.^10, 11^ Moreover, the MTBRs not only bind to microtubules but also to acidic membranes,^12, 13^ implicating direct binding to the inner leaflet of the plasma membrane. The fact that both microtubules^14^ and acidic membranes have highly negatively charged surfaces raises the prospect of mimicry in their binding by tau. This prospect is supported by two other observations. First, the MTBRs bind DNA^15^ and RNA.^16^ which also have highly negatively charged surfaces. Second, tau forms aggregates at acidic membrane surfaces^17, 18^ and condensates known as “hot spots” at the plasma membrane,^19^ similar to tau condensate formation over microtubules.^5^ A common mechanism for the enhancement of aggregation or condensation may be that the negatively charged surfaces serve to neutralize and concentrate tau.^20^

The atomic structure of the tau-microtubule complex was recently determined by cryo-electron microscopy (cryo-EM).^21^ In contrast, information about the complex between tau and acidic membranes is much more limited, due to the highly dynamic nature of this complex. Solid-state NMR has revealed the involvement of Lys residues in the binding of tau K19, a construct comprising MTBRs R1, R3, and R4 (Fig. 1A), to DMPC/DMPS membranes, and the participation of a subset of Val residues in helix formation at the DMPS membrane surface.^12^ Electron paramagnetic resonance (EPR) data have further defined the residues forming amphipathic helices (Fig. 1A, B) when tau K19 is bound to POPC/POPS membranes.^13^ Molecular dynamics (MD) simulations have much to offer in characterizing atomic details of IDP binding to membranes, as demonstrated in our recent studies.^22–24^

**Figure 1.**
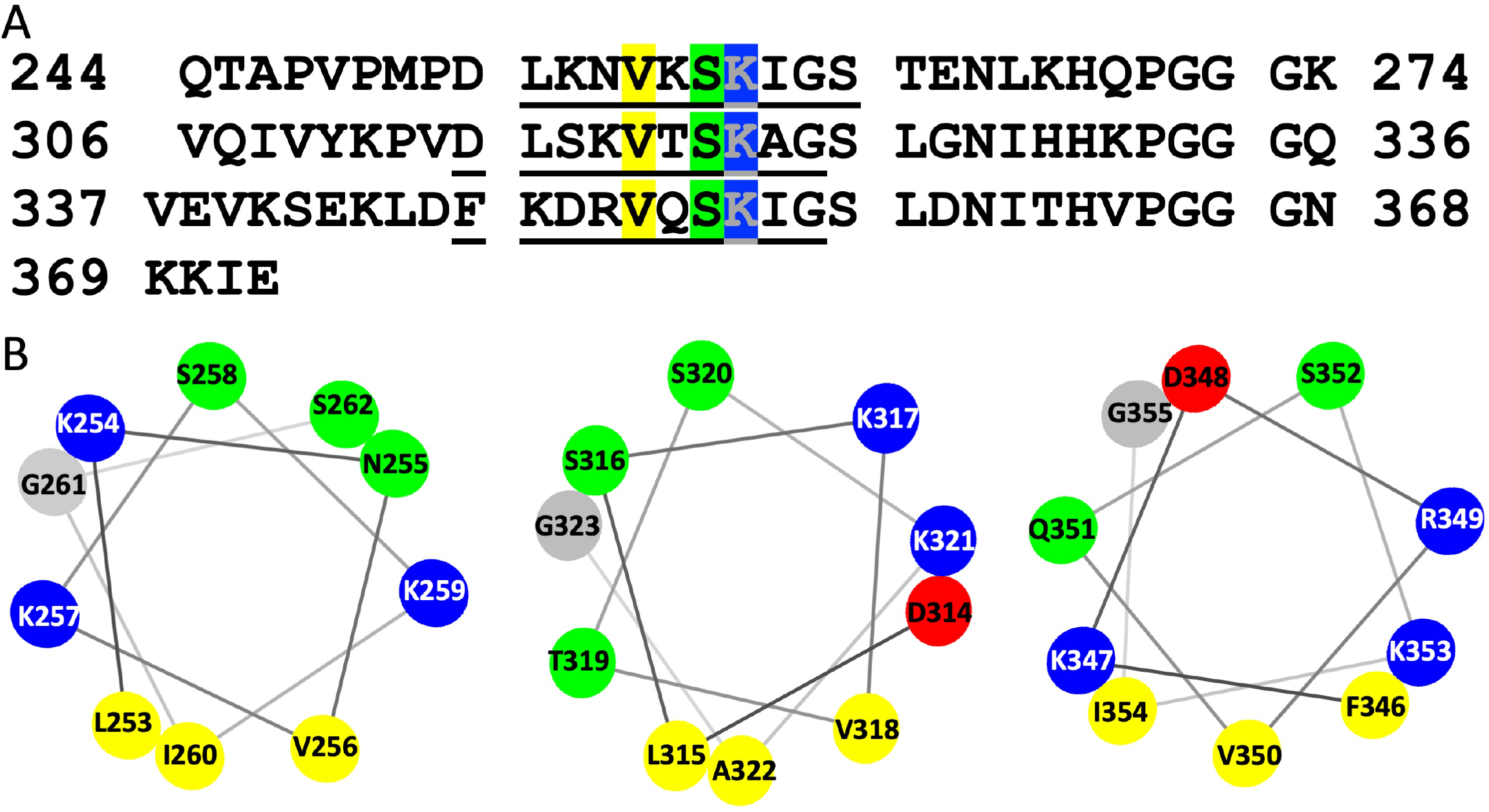
Amino-acid sequence and amphipathic helices of tau K19. (A) Sequence separated into three repeats (first three lines), with helical residues underlined and conserved residues in the helices shaded in different colors. (B) Helical wheel representation for residues 253-262, 314-323, and 346-355. Nonpolar, polar, acidic, basic, and Gly residues are shown in yellow, green, red, blue, and grey, respectively.

Here we report MD simulation results of tau K19 bound to POPC/POPS membranes. Our MD simulations show that three amphipathic helices, one in each repeat, stably bind to the membrane. In each amphipathic helix, a Lys that is conserved among the MTBRs, along with other charged and polar sidechains, interacts with lipid headgroups, while a conserved Val along with two other nonpolar sidechains inserts into the hydrophobic region of the membrane. Interestingly, we note these conserved residues also form similar interactions with microtubules. We propose that this partial mimicry facilitates both the membrane anchoring of microtubules by tau and the transfer of tau from membranes to microtubules.

## Results

### An Amphipathic Helix in Each Repeat of Tau K19 Stably Binds to POPC/POPS Membranes

The sequence of tau K19, separated into three repeats, is shown in Fig. 1A. Based on published EPR data^13^ and on the amphipathic sequence pattern (Fig. 1B), we modeled 10 residues in each of the three repeats in tau K19 as a helix and placed it at the hydrophilic-hydrophobic interface of POPC/POPS (1:1) membranes. Apart from modest fraying in the helix in repeat 1 (helix 1), the helices are well maintained in four 600-ns long replicate simulations. As illustrated by a snapshot shown in Fig. 2, the three helices also stably bind to the membrane.

**Figure 2.**
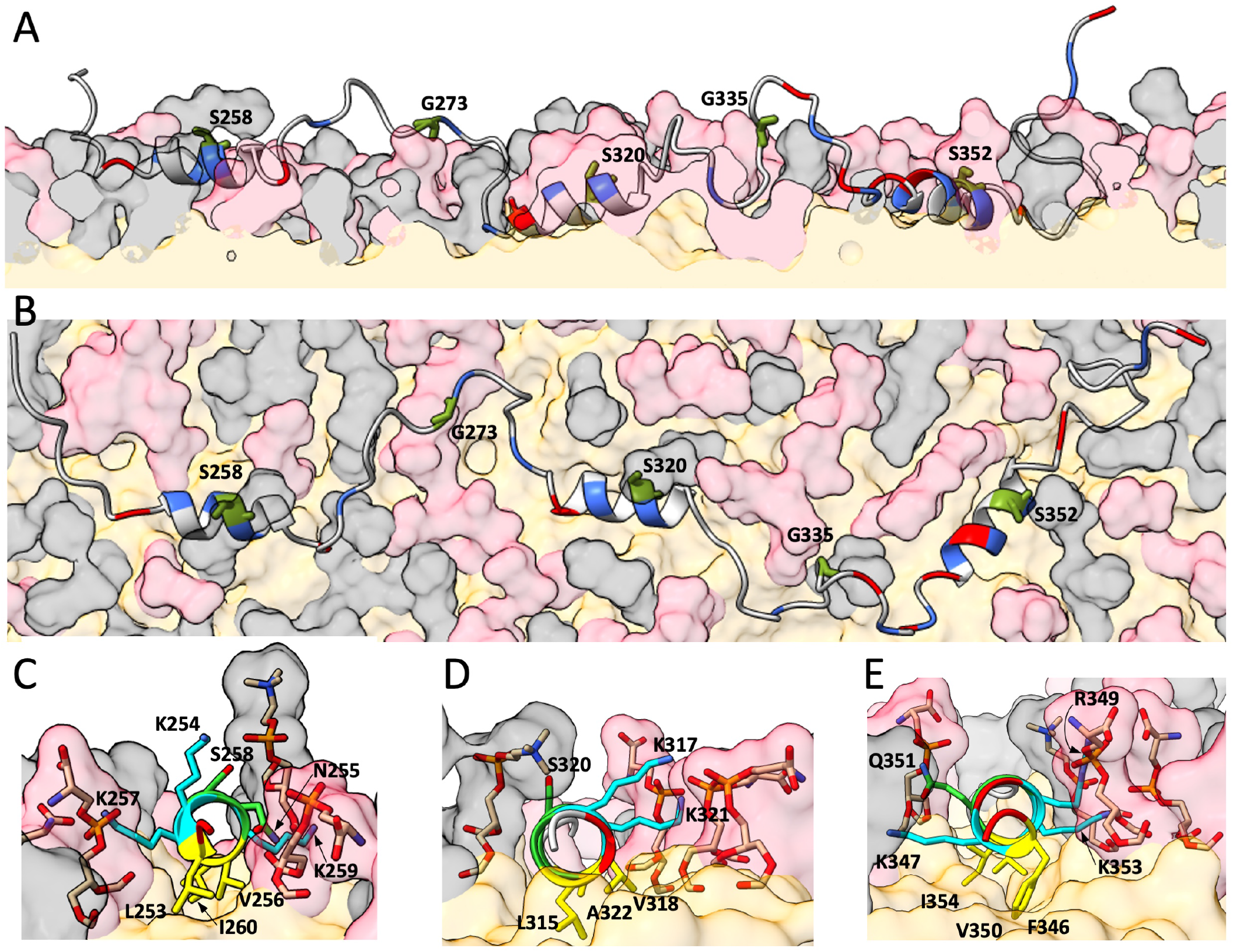
A snapshot of tau K19 bound to a POPC/POPS membrane in MD simulations. (A-B) Side and top views. Acidic and basic residues are colored in red and blue, respectively. Residues where spin labels were introduced are in olive green and additionally shown as sticks. Lipids are shown as surface representation, with POPC and POPS headgroups and acyl chains in grey, pink, and orange, respectively. (C-E) Images of membrane-bound helices 1, 3, and 4. Nonpolar, polar, acidic, and basic residues in the helices are shown in yellow, green, red, and cyan, respectively. The nonpolar, basic, and notable polar sidechains as well as lipid headgroups in contact with protein sidechains are additionally shown as sticks, with O and N atoms in red and blue, respectively.

As the first validation of the MD simulations against experimental data, we calculated the distributions of distances between spin labels (MTSSL) at seven pairs of positions, and compared them with the counterparts obtained using double electron-electron resonance (DEER).^13^ The seven pairs of distances are between Ser residues at a conserved position (Ser258, Ser320, and S352; see Fig. 1A) in the three repeats, or between a Gly residue at the C-terminus of repeat 1 or 3 (Gly273 or Gly335) and an upstream or downstream neighboring conserved Ser. The three Ser residues are at the top of the amphipathic helices while the two Gly residues are near the middle of the linker between two adjacent helices (Fig. 2A, B). Figure 3 shows that the distance distributions calculated from the MD simulations overlap well with the DEER results. The distance distributions are broad, partly due to the flexibility of the spin labels. They peak around 50 Å for a pair of Ser residues in two neighboring helices (helices 1 and 3 or helices 3 and 4) and around 30 Å between a terminal Gly and a neighboring conserved Ser. For the seventh pair, between the Ser residues in helices 1 and 4, the DEER data only placed a lower bound of 60 Å for the peak distance; the peak distance, 82 Å, from the MD simulations is consistent with this bound.

**Figure 3.**
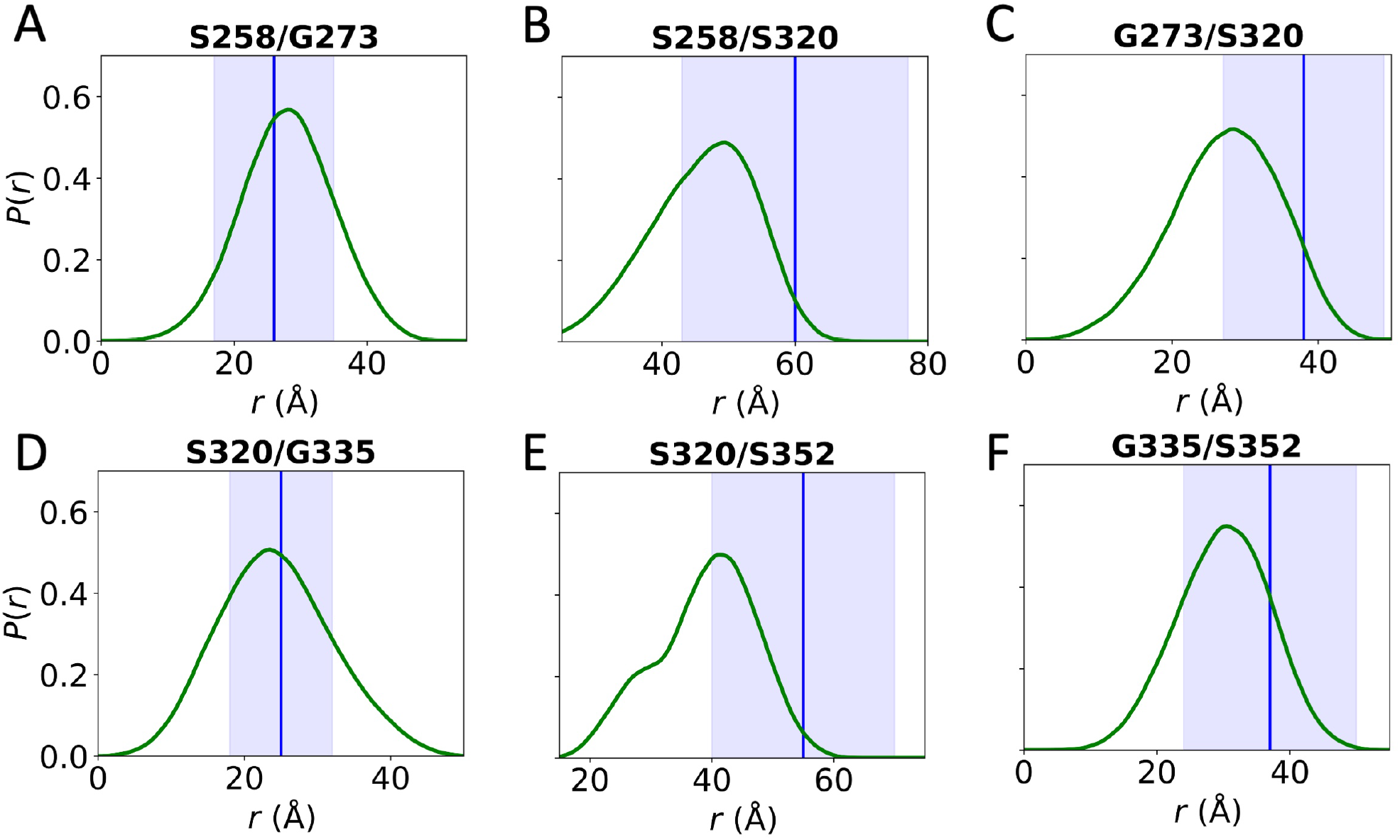
Distributions of interhelical and helix-linker distances. (A-F) Distributions for the distances between spin labels at the indicated pair of residues. MD results are shown as green curves; experimental peak distances and standard deviations are shown by a blue vertical line and shading.

### Interactions Mediating Tau K19-Membrane Binding Show Both Similarities and Distinctions Among the Three Repeats

To identify the residues that mediate tau K19-membrane binding, we used two complementary measures. The first is *Z*_tip_, the displacement of the sidechain heavy tip atom of each residue from the membrane surface, defined as the mean plane of the lipid phosphorus atoms. The second is the membrane contact probability, calculated as the fraction of MD snapshots in which a residue is in contact with the membrane (i.e., with at least one pair of heavy atoms within 3.5 Å).

The *Z*_tip_ results are presented in Fig. 4. Helix 1 is buried relatively superficially in the membrane, with mean *Z*_tip_ values ranging from 0 Å for the nonpolar sidechains (Leu253, Val256, and Ile260) at the bottom to 6 Å for polar and charged sidechains (Lys254, Asn255, Ser258, and Ser262) at the top. The oscillation in the mean *Z*_tip_ values is due to the positioning of the residues around the helix (see Fig. 1B). Helix 3 is more deeply buried, with mean *Z*_tip_ values around −5 Å for the nonpolar sidechains (Leu315, Val318, and Ala322) at the bottom and around 2 Å for sidechains (Lys317 and Ser320) at the top. This helix is tilted so the burial is deeper at the N-terminus. Helix 4 is buried at a similar depth as helix 3, but tilted in the oppositely direction, such that the mean *Z*_tip_ values of the nonpolar sidechains (Phe346, Val350, and Ile354) progressively decrease, reaching −7 Å at Ile354. At the top of this helix, Asp348 and Ser352 have mean *Z*_tip_ values around 1 Å. In addition, the C-terminal extension of five residues, with sequence SLDNI, has a tendency to form 310 helix (Fig. S1), and is also positioned near the membrane hydrophobic-hydrophilic interface. Indeed, the two nonpolar sidechains, Leu357 and Ile360 are buried as deep as the third nonpolar residue, Ile354, within helix 4. The middle of each interhelical linker as well as the two termini of tau K19 are well above the membrane surface.

**Figure 4.**
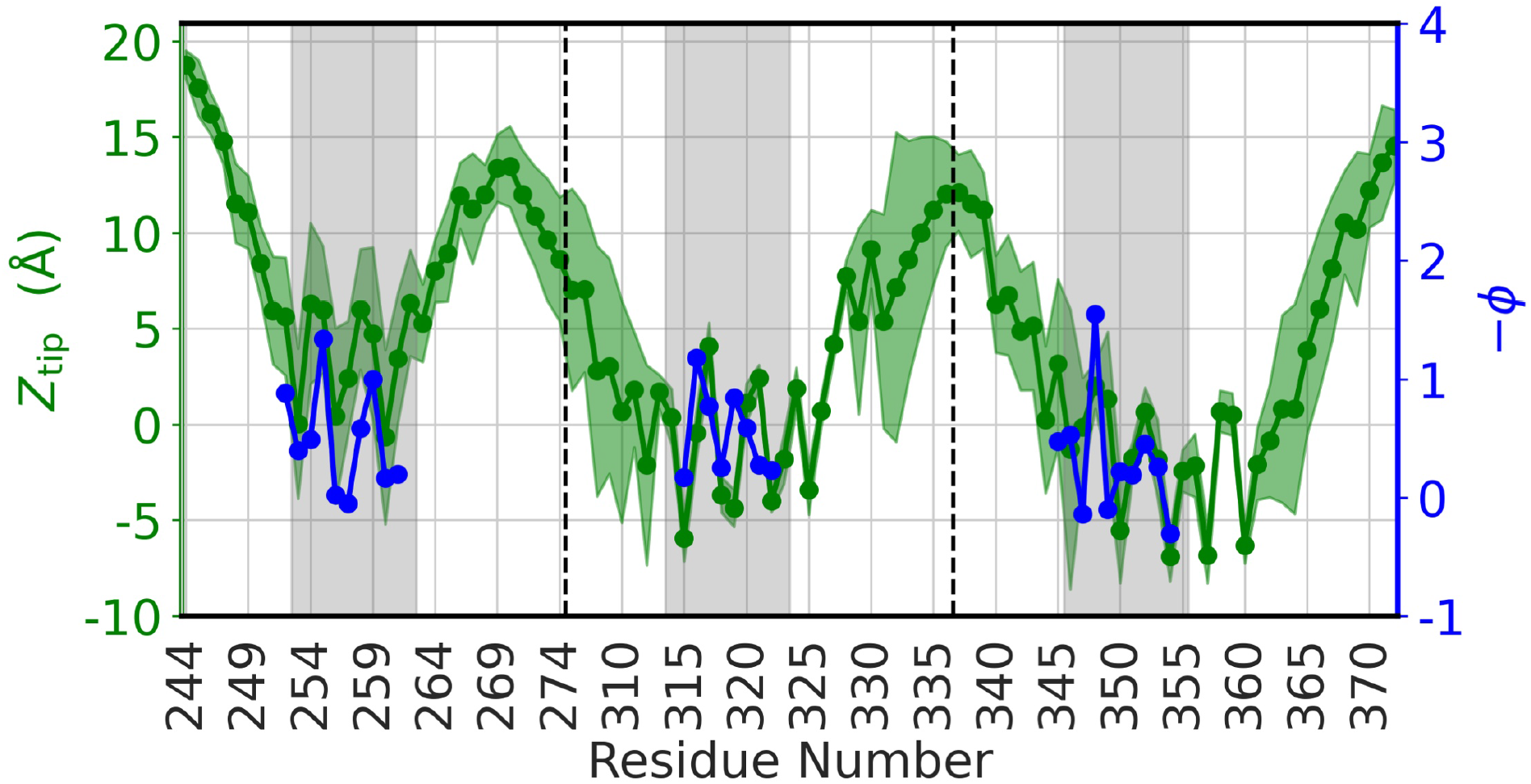
*Z*_tip_ distances from MD simulations and depth parameter −*Φ* from EPR. Mean values and standard deviations of *Z*_tip_ distances are shown as green circles and shading, and reference the left axis; −*Φ* values are shown as blue circles and reference the right axis. Vertical lines demarcate each repeat region and grey shading covers the helical region in each repeat.

Figure 2A provides illustration for the different extents of burial between helix 1 and helices 3 and 4, the opposite tilting of helices 3 and 4, and the burial of the C-terminal extension of helix 4. Figure 2C-E further shows that, in helices 3 and 4, the nonpolar sidechains cross a flat hydrophilic-hydrophobic interface into the hydrophobic region of the membrane, whereas in helix 1 it looks as if local acyl chains move up to surround the nonpolar sidechains. Note that the second of the three nonpolar residues in each helix is a conserved Val (see Fig. 1A).

The *Z*_tip_ results are validated by EPR data obtained from accessibility to spin labels.^13^ The same oscillation patterns of mean *Z*_tip_ values in the three helical segments are exhibited by the experimental depth parameter −*Φ*. Low mean *Z*_tip_ values for the three nonpolar sidechains in each helix are matched by low *–Φ* values, while high mean *Z*_tip_ values for the sidechains on top of the helices, including Asn255 and Ser258 in helix 1, Ser320 in helix 2, and Asp348 and Ser352, are matched by high –*Φ* values. The shallower burial of helix 1 also seems to be supported by the accessibility data: when the –*Φ* values for each helical region were fit to a sine function, the resulting function for helix 1 reached a higher level than those for helices 3 and 4.

The membrane contact probabilities corroborate the *Z*_tip_ results but also provide additional information (Fig. 5A). As expected, the three nonpolar sidechains have the highest or close to the highest membrane contact probabilities, which are 90% for helix 1 and 100% for helices 3 and 4. In contrast, sidechains at the top of each helix have lower membrane contact probabilities, dipping below 50% for Lys254, Asn255, and Ser258 in helix 1, Lys317 in helix 3, and Asp348 and Ser352 in helix 4.

**Figure 5.**
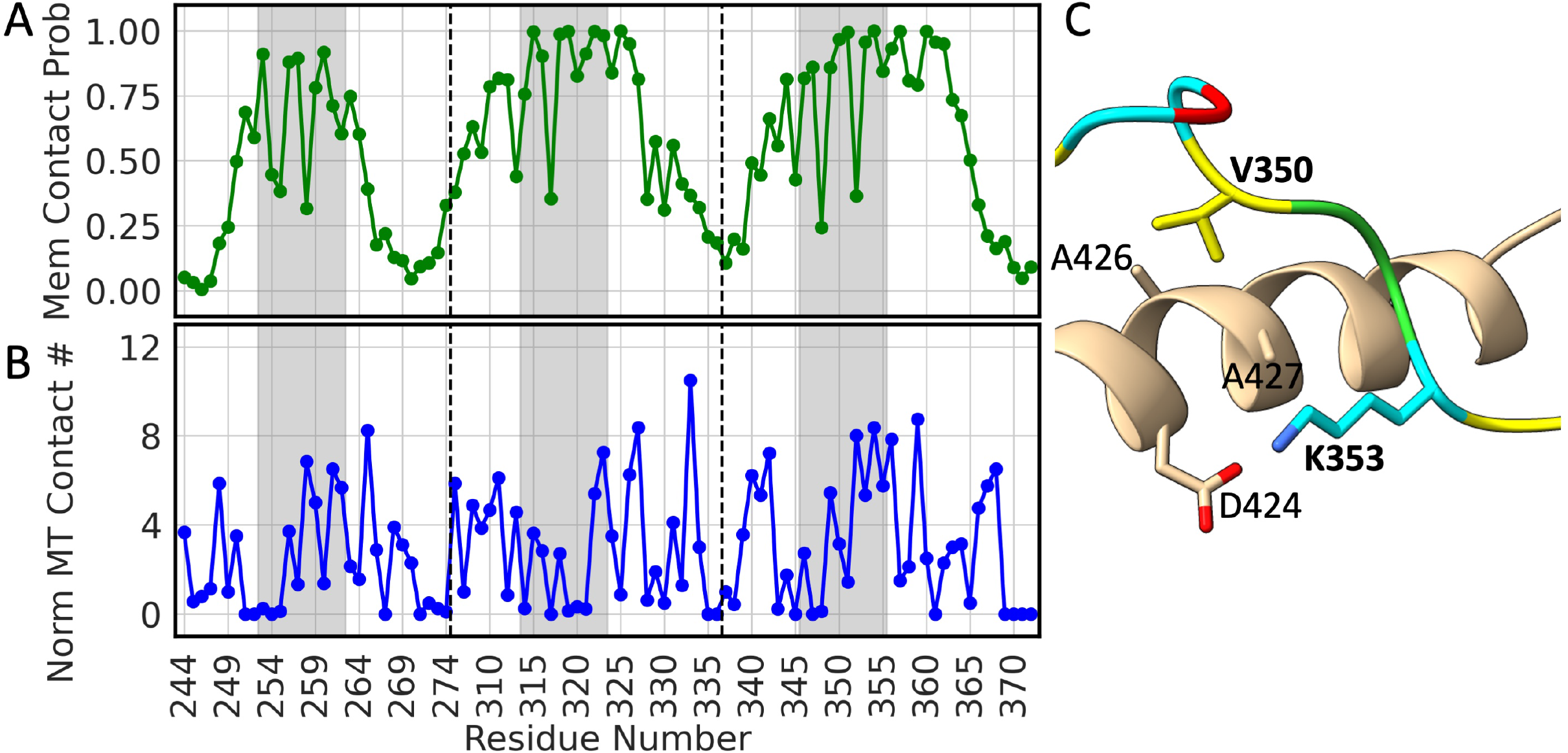
Membrane and microtubule contact properties of tau K19. (A) Membrane contact probabilities of tau residues. (B) Normalized microtubule contact numbers of tau residues. The number of microtubule residues in contact with a tau residue was calculated from Protein Data Bank entry 7PQC and then normalized by the number of heavy atoms in that residue. A low contact number for Lys321 can be attributed to the particular model deposited in 7PQC (see also Fig. S2D). (C) Image of tau-microtubule interaction in repeat 4, highlighting the roles of a conserved Val and a conserved Lys. Nonpolar, polar, acidic, and basic residues of tau are shown in yellow, green, red, and cyan, respectively; α-tubulin is shown in tan. Residue labels for tau are in bold.

In addition to the insertion of the nonpolar sidechains into the hydrophobic region, membrane binding of each amphipathic helix is also stabilized by electrostatic interactions of charged and polar sidechains, located on the sides of the helix, with lipid headgroups. Among these is a conserved Lys (see Fig. 1A, B), three positions downstream of the conserved Val. These Lys residues, at 259, 321, and 353, all have membrane contact probabilities close to the maxima in their respective helices. As shown in Fig. 2C-E, their sidechains project sideways, with the nonpolar portion atop the hydrophobic region and the terminal amine tipping upward into the headgroup region. In both helices 1 and 4, a Lys (at 257 and 347) occupies a similar location as the conserved Lys but on the opposite side of the helix. Unique to helix 4, an Arg residue (at 349) is positioned just above the conserved Lys, with its guonidino group pointing upward to form extensive interactions with lipid headgroups. In the same helix, Gln351 sits opposite of Arg349, with its amide reaching into the headgroup region. Additional residues with high membrane contact probabilities are two residues with short polar sidechains, Ser316 and Thr329, in helix 3 and two Gly residues, 323 in helix 3 and 355 in helix 4.

In short, the three helices have three conserved residues, each representing a different facet of an amphipathic helix. The conserved Ser sits on top of the helix and participates minimally in lipid interactions; the conserved Val is at the bottom of the helix and reaches into the hydrophobic region of the membrane; the conserved Lys projects sideways and forms electrostatic interactions with lipid headgroups. These roles of Val and Lys residues are supported by solid-state NMR data.^12^ Specifically, ^13^C-^13^C correlation spectra of tau K19 bound to DMPC membranes showed that a subset of Val residues participate in helix formation. We now suggest that this subset consists of the three Val residues (out of a total of 10 Val residues) at the conserved position in the three amphipathic helices. Furthermore, ^1^H-^13^C heteronuclear correlation spectra showed the involvement of Lys residues in the membrane binding of tau K19. Our simulation results document significant contributions of Lys residues, at both the conserved position and other nonconserved positions, to membrane binding.

There are also significant differences among the three helices in membrane interactions. The most notable is the much shallower burial of helix 1 relative to helices 3 and 4. One reason for this difference is the high number of polar and charged residues at the top of helix 1. This number is five for helix 1, compared to three for helix 3 or 4. These helix-top sidechains have limited ability to interact with lipids thus contribute relatively little to membrane biding. A second reason is flanking residues. We have already noted the membrane insertion of the C-terminal extension of helix 4. According to the membrane contact data in Fig. 5A, no flanking residues of helix 1 have > 75% membrane contact probabilities, but there are seven such residues for helix 3 and eight such residues for helix 4. The flanking residues may also be responsible for the opposite tilts of helices 3 and 4. The membrane insertion of the C-terminal extension may explain the deeper burial at the C-terminus of helix 4. For helix 3, only the first part of the C-terminal extension is buried; instead, the tilt of this helix appears to be driven by Lys311 in the N-terminal extension, which is well positioned in the headgroup region (see Fig. 2A).

### Partial Mimicry Between Microtubule Binding and Membrane Binding

The microtubule-bound structure of a tau construct comprising the four MTBRs plus flanking regions (residues 202 to 395) was recently determined by cryo-EM (Protein Data Bank entry 7PQC).^21^ The MTBRs are stretched into a linear conformation lying atop the microtubule surface (Fig. S2A). At first glance, there is very little resemblance between the microtubule-bound tau and the membrane-bound tau (Fig. 2B), which forms a helix in each repeat. Closer inspections, however, revealed interesting mimicry. To assess the similarities and differences of tau-microtubule interactions to tau-membrane interactions, we calculated microtubule contact numbers of tau (i.e., number of microtubule residues within 5 Å of a tau residue) in the cryo-EM structure (Fig. 5B). We also highlight some of the tau-microtubule interactions in Figs. 5C and S2B-F.

The most striking similarity involves the conserved Lys and Val residues. As shown in Fig. 2C-E, when tau is bound to acidic membranes, the Lys sidechains form electrostatic interactions with lipid headgroups while the Val sidechains (the second of three nonpolar resides in each amphipathic helix) insert into the hydrophobic region. On the microtubule surface, the conserved Lys residues form a salt bridge with a-tubulin Asp424, located on the side of an a-helix (Figs. 5C and S2B-D), while the conserved Val residues form hydrophobic interactions with a pair of Ala residues (426 and 427) at the top of the same a-helix (see also Fig. S2E). The high membrane contact probabilities of these conserved residues are matched by the high microtubule contact numbers per tau heavy atom (Fig. 5A, B). The third nonpolar sidechain (Ile260, Ala322, or Ile354 in tau K19), previously found in the membrane-bound amphipathic helix, also projects into a hydrophobic pocket in the microtubule-bound structure, with the fit being the best in repeat 4 (shown in Fig. S2E). On the other hand, the residues facing the first nonpolar sidechain (Leu253, Leu315, or Phe346 in tau K19) are all polar or charged. Correspondingly the microtubule contact numbers of these nonpolar residues are relatively low.

The second similarity involves some of the residues at the top of the amphipathic helices in the membrane-bound state and have < 50% membrane contact probabilities. Of these, Lys254 and Asn255 in repeat 1, Lys317 in repeat 3, and Asp348 in repeat 4 all have their sidechains projected away from microtubules, and therefore have minimal microtubule contact numbers. That is, these residues participate minimally in both membrane and microtubule binding.

The third similarity lies in the lower level of repeat 1 in engaging with the target surfaces when compared the downstream repeats. As reported above, helix 1 has a shallower burial into the membrane relative to helices 3 and 4. Brotzakis et al.^21^ also noted that repeat 1 is less stably bound to microtubules than repeats 2 to 4. The differences between repeats can be quantified by membrane contact probabilities and microtubule contact numbers. The number of residues with > 75% membrane contact probabilities is 5 for repeat 1 but goes up to 16 each for repeats 3 and 4. Likewise, the number of tau residues with > 4 microtubule contact numbers per heavy atom is 6 for repeat 1 but is approximately doubled, to 11 and 13, respectively, for repeats 3 and 4. For repeat 4, we have already mentioned that the hydrophobic fit of the third nonpolar sidechain is the best. In addition, Arg349, which as part of the fourth amphipathic helix forms extensive electrostatic interactions with lipid headgroups (Fig. 2E), also forms a salt bridge with Glu437 in the C-terminal tail of a neighboring β-tubulin (Fig. S2F).

Tau’s interactions with microtubules also have noticeable differences from the counterparts with membranes. The fact that the first nonpolar sidechain faces a polar environment on the microtubule surface, as opposed to the hydrophobic region within the membrane, has been stated above. In addition, the conserved Ser residues (at 258 and 352) at the top of a tau amphipathic helix have relatively low membrane contact probabilities, but on the microtubule surface they hydrogen bond to Glu434 in one a-tubulin chain and Asp431 in another a-tubulin chain, respectively. Lastly, a partial difference between membrane and microtubule binding is observed on a conserved Asn (at 265, 327, or 359). The conserved Asn residues in repeats 3 and 4 participate in membrane binding as part of the C-terminal extension of an amphipathic helix, but the counterpart in repeat 1 is above the membrane surface. In comparison, the conserved Asn residues in all the repeats participate in microtubule binding, forming a hydrogen bond with β-tubulin Lys392 and/or an amino-π interaction with β-tubulin Phe389.

## Discussion

Using MD simulations, we have characterized the atomic details of tau K19 binding to acidic membranes. An amphipathic helix is formed in each of the three repeats to anchor the membrane binding. These helices have three conserved residues, each representing a different facet of membrane interactions. A Val sidechain at the bottom of the helix inserts into the hydrophobic region whereas a Lys sidechain projects sideways into the headgroup region. In contrast, a Ser at the top of the helix minimally participate in membrane interactions. Other residues in the amphipathic helices as well as N- and C-terminal extensions provide distinctions to the three repeats in membrane binding stability. In particular, helix 1 is less stably bound than the downstream helices. Previous experimental data provide validation of the simulation results, including the distances between the helices, the burial depths of helical regions, and the roles of Val and Lys residues in membrane binding.

Importantly, we can recognize mimicry between tau-membrane binding characterized here and tau-microtubule binding revealed by cryo-EM. This mimicry is not as one would naively expect, where basic residues in the tau MTBRs would attach to negatively charged surfaces presented by acidic membranes or microtubules. Rather, both charged and nonpolar residues of tau participate in target binding, but in ways different for membranes and microtubules. Both lipid membranes and microtubules present environments that are highly heterogeneous, but in different manners. In acidic membranes, the headgroups and acyl chains separate into charged and hydrophobic layers. The formation of an amphipathic helix allows the nonpolar sidechains (e.g., Val) to insert into the acyl chains and the basic sidechains (e.g., Lys) to project into the headgroup region. Microtubules, on the other hand, feature charged and nonpolar surface patches right next to each other. On the microtubule surface, tau MTBRs stretch into a linear conformation, with nonpolar sidechains including the conserved Val fit into hydrophobic pockets whereas adjacent basic sidechains including the conserved Lys form salt bridges with acidic residues of tubulins.

The membrane binding of tau MTBRs and its partial mimicry of microtubule binding have significant implications for the functions and disease linkage of tau. A substantial portion of dephosphorylated tau is associated with membranes.^11^ Hyperphosphorylation results in the release of tau from membranes, thereby promoting the formation of neurofibrillary tangles from cytosolic tau.^25^ Membrane association of tau may occur indirectly by binding to membrane-bound proteins such as annexins^8^ and Fyn,^9^ or directly through the MTBRs as characterized here and previously^12, 13^ (Fig. 6). Membrane-associated tau may anchor microtubules to membranes,^8^ and may also further concentrate to form “hot spots”.^19^ When a microtubule comes into contact with tau hot spots at the membrane surface, multiple copies of tau may transfer onto the microtubule surface. The partial mimicry between tau binding to the two target surfaces may facilitate both the membrane anchoring of microtubules by tau and the transfer of tau from membranes to microtubules.

**Figure 6.**
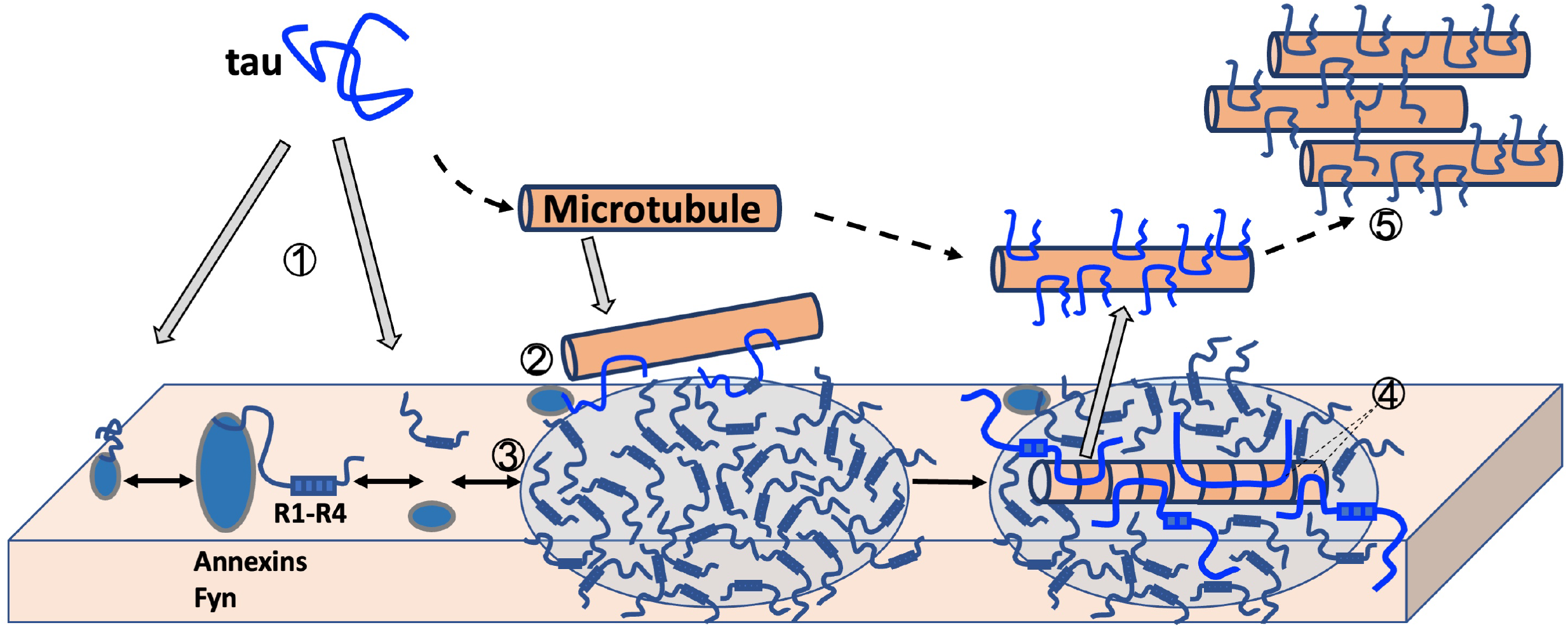
Functional steps initiated by membrane association of tau. The illustrated steps are: (1) membrane association by indirect (via binding to membrane-bound proteins) or direct (via MTBRs) routes; (2) membrane anchoring of microtubules by tau; (3) concentration of tau at the membrane surface; (4) membrane-to-microtubule transfer of tau; and (5) condensate formation of microtubule-bound tau. Due to binding mimicry, a single tau molecule can have some MTBRs bound to the membrane and other MTBRs bound to a microtubule. This simultaneous binding provides an additional mechanism for the membrane anchoring of tau and serves as an intermediate in the transfer of tau from membranes to microtubules.

## Computational Methods

### Molecular Dynamics Simulations

MD simulations of membrane-bound tau K19 were ran using AMBER18^26^ with the ff14SB^27^ force field for protein, lipid17^28^ for membrane, and TIP4P-D^29^ for water, as in our previous studies.^22–24^ To construct an initial model for tau K19, the helical segments were built using pymol (https://pymol.org/) while the linkers and disordered tails were generated using TRADES.^30^ The initial depth of the helices was adjusted to have complete burial of the nonpolar sidechains in the hydrophobic region of a membrane. CHARMM-GUI^31^ was then used to insert into a membrane without changing the depth and solvate with water and 10 mM NaCl. The lipid composition was POPC/POPS at a 1:1 ratio, chose to match with the experimental condition of Georgieva et al.^13^ The number of lipids was 330 in the upper leaflet; 25 extra POPC lipids were added to the lower leaflet in order to maintain parity in surface area between the two leaflets. The dimensions of the simulation box were 151 Å × 151 Å × 117 Å; the total number of atoms was 303970.

The system was first prepared in short MD simulations in NAMD. Following 10000 steps of steepest-decent energy minimization, 675 ps of equilibration was split across six separate steps, where restraints were successively reduced to 0 as the simulation shifted from constant NVT to constant NPT (298 K and 1 atm pressure) and the timestep changed from 1 fs to 2 fs. The system was then split into four replicates with different random seeds, each running 600 ns in AMBER at constant NPT with a timestep of 2 fs on GPUs using *pmemd.cuda*^32^ The SHAKE algorithm^33^ was used to restrain bond lengths involving hydrogens. The particle mesh Ewald method^34^ was used to treat long-range electrostatic interactions. The cutoff for nonbonded interactions was 12.0 Å. Temperature was regulated by the Langevin thermostat^35^ with a 1.0 ps^−1^ dampening constant and pressure was regulated by the Berendsen barostat^36^ with a coupling constant of 0.5 ps. Frames were saved every 20 ps for analysis, unless otherwise indicated.

### Data Analysis

Secondary structures were calculated using the DSSP algorithm implemented in CPPTRAJ with the secstruct command.^37^ Distance distributions emulating DEER data were calculated using the DEER-PREdict software,^38^ averaged over frames saved every 1 ns. *Z*_tip_ distances were calculated by subtracting the mean *Z* coordinate of the lipid phosphorous atoms in the upper leaflet in a single frame from the *Z* coordinate of the sidechain heavy tip atom of each residue. For membrane contact probabilities, a contact was defined when a heavy atom of the protein residue came within 3.5 Å of any heavy atom of the membrane. The fraction of frames where at least one membrane contact was formed with a given residue was calculated as the membrane contact probability of that residue. The microtubule contact number of each tau residue was calculated from the cryo-EM structure (Protein Data Bank entry 7PQC) using a 5 Å cutoff between heavy atoms, and then normalized by the number of heavy atoms in that tau residue.

## Acknowledgments

This work was supported by National Institutes of Health Grant R35 GM118091.

## Supporting Information Available

Two supporting figures, presenting secondary structures of membrane-bound tau K19 in MD simulations and interactions of tau with microtubules in a cryo-EM structure.

## TOC Graphic

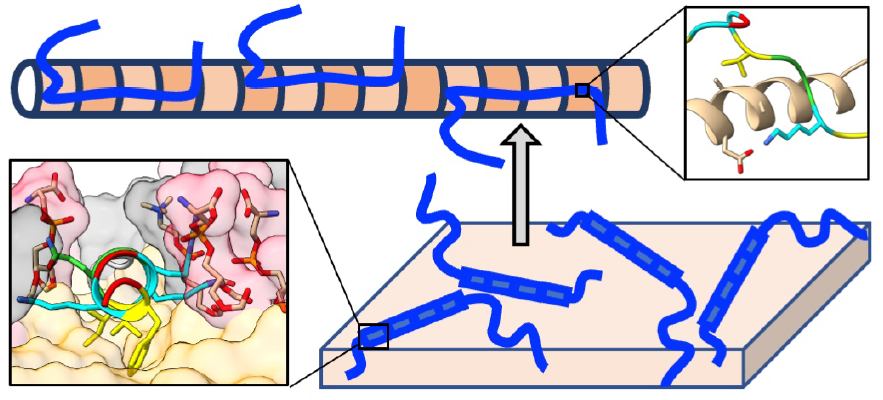

