## Supporting Figures for "Partial Mimicry of the Microtubule Binding of Tau by Its Membrane Binding"

Matthew MacAinsh[1], and Huan-Xiang Zhou[1,2,*]
[1]Department of Chemistry and [2]Department of Physics, University of Illinois at Chicago, Chicago, IL 60607, USA

*Supporting Information*

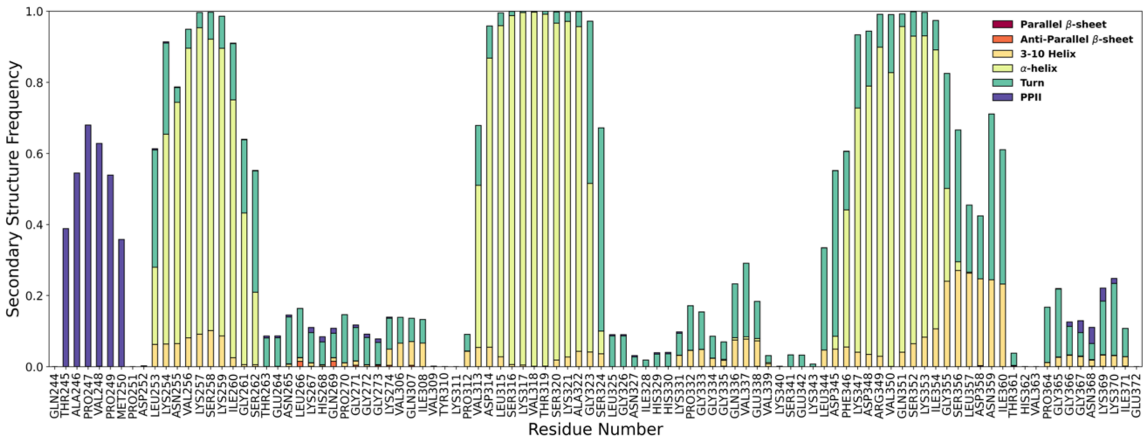


Figure S1. Secondary structure propensities of membrane-bound tau K19 in MD simulations.

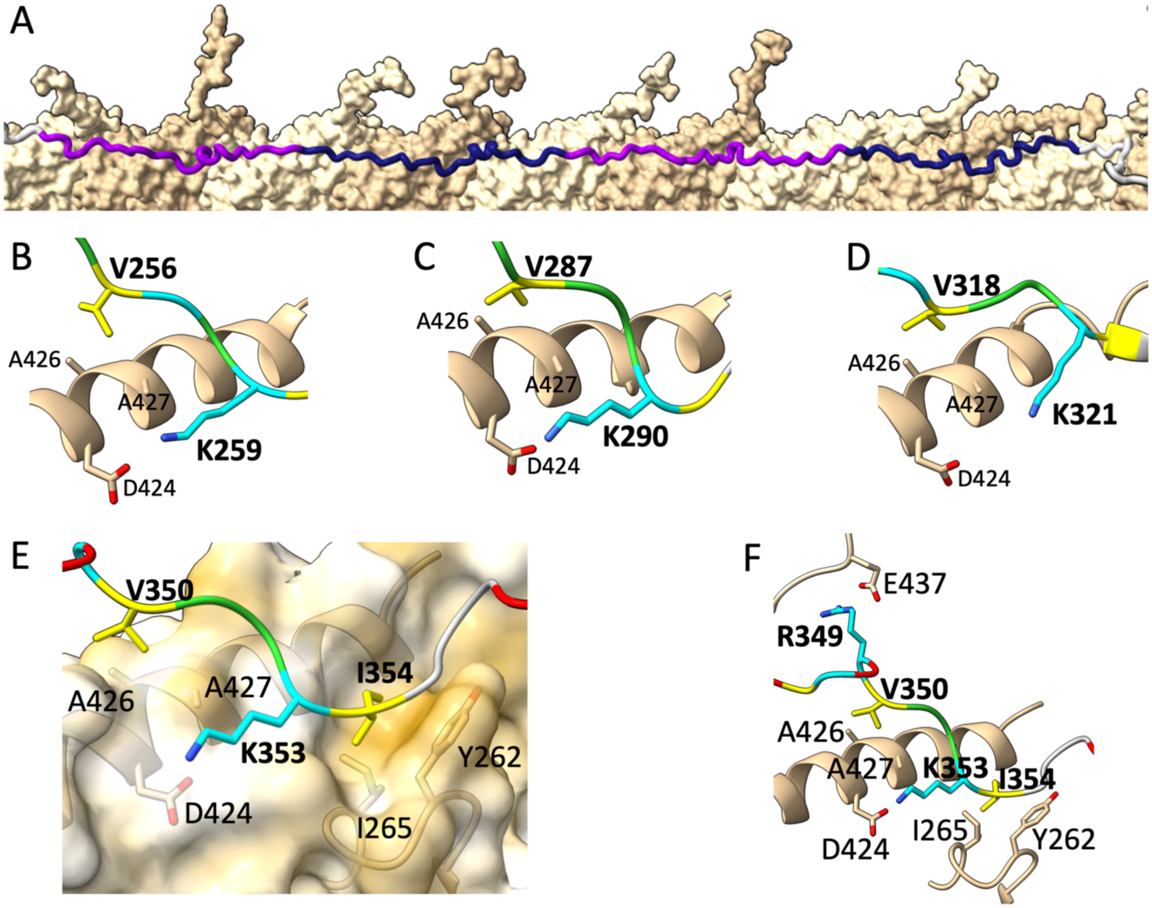


Figure S2. Tau-microtubule interactions in Protein Data Bank entry 7PQC. (A) Linear conformation of tau on the microtubule surface. α- and β-tubulin are in darker and lighter tan colors, respectively; tau MTBRs are in alternative light or dark purple colors, with flanking regions in gray. (B-C) Conserved Val and Lys residues of tau in repeats 1, 2, and 3 interacting with a microtubule. (E) A portion of repeat 4 on top of the microtubule surface, shown in a hydrophobicity scale with hydrophobic patches in gold and hydrophilic patches in white. (F) View of repeat 4 interacting with a microtubule, expanded to include both the conserved Val and Lys and the additional Arg349, which forms a salt bridge with β-tubulin Glu437. Nonpolar, polar, acidic, and basic residues of tau are in yellow, green, red, and cyan, respectively; α- and β-tubulin are in tan. Residue labels for tau are in bold.
